# Metagenomic analysis of the effects of European bison *Bison bonasus* (Linnaeus, 1758) presence on soil community structure and function in West Blean and Thornden Woods, Kent

**DOI:** 10.64898/2026.08.19.745704

**Authors:** Chonghan Xu, Hester Van Schalkwyk, Owain Powell, Carla Gustave, Lawrence Ball, Kora Ross, Edward Murray, Heidi Aguirregoicoa, Hannah Mackins, Kirsty Swinnerton, Thomas J. Creedy, Laura Sivess, Jumayma Jones, Kath Castillo, Rosie Bleet, Silvia Salatino, Yuen-Ting Chan Mendis, Raju Misra, Darren Chooneea, Pedro Lebre, Hermine Mkrtchyan, Piotr Cuber

## Abstract

The reintroduction of extinct or endangered species to restore ecosystem function is an essential aspect of rewilding. The Wilder Blean Project at West Blean and Thornden Woods in Canterbury, UK, is committed to rewilding natural processes and enhancing biodiversity in one of England’s oldest and largest areas of ancient woodland. The introduction of European bison (*Bison bonasus*) is an important part of the project. However, how the reintroduction of large herbivores influences local biodiversity and ecosystem functions during the early stages of rewilding remains poorly understood. Soil samples were collected from the same sampling sites before and two years after bison were reintroduced and profiled by metagenomic sequencing using Oxford Nanopore Technologies sequencing platforms. The results showed that the alpha diversity of soil organisms did not change significantly before and after the introduction of European bison, while beta diversity showed modest shifts in community composition. The relative abundance of some nitrogen-fixing and photosynthetic microbial genera showed declines in the 2024 Bison Area, while the mycorrhizal fungus genus Rhizophagus was significantly less abundant than in the 2024 Control Area. Despite relatively stable taxonomic diversity, functional composition differed significantly between the 2022 and 2024 Bison areas and among the 2024 rewilding treatments, revealing a decoupling between taxonomic diversity and functional composition. Amino acid synthesis pathways and carbon metabolism pathways were significantly enriched. These findings highlight the potential of long-read Oxford Nanopore metagenomics to reveal functional shifts that may not be apparent from taxonomic diversity alone. Although these early-stage responses cannot yet predict long-term rewilding trajectories, continued longitudinal monitoring integrating microbial, soil physicochemical, and ecosystem-level measurements will be essential to determine the persistence and ecological significance of these functional shifts.

## 1. Introduction

A growing body of literature emphasises the need for a process-oriented approach to ecosystem restoration in our rapidly changing world, rather than one aimed at maintaining or restoring species with a predefined composition (Valiente-Banuet et al., 2014; Higgs et al., 2018). Rewilding is one such restoration approach. This strategy aims to restore self-sustaining complex ecosystems while minimising or phasing out human intervention (Lorimer et al., 2015; Jepson, 2015; Fernández et al., 2017). Trophic level rewilding is an important aspect of rewilding that promotes the reintroduction of key missing species such as large carnivores and large herbivores (Perino et al., 2019). The selective loss of top predators and large herbivores may lead to trophic cascade effects and a greater susceptibility to ecosystem collapse (Beschta & Ripple, 2009; Ripple et al., 2015; Letnic et al., 2009). Trophic rewilding also promotes the use of functional substitutes, i.e., the introduction of non-native species as ecological replacements for species that became extinct centuries or millennia ago (Jørgensen, 2015; Svenning et al., 2016; Fernández et al., 2017). The European bison (*Bison bonasus* L., 1758) is the key stone species of the burgeoning rewilding movement (Lord et al., 2020; Kuemmerle et al., 2011). As the largest herbivorous mammal in Europe, the species was recognised as extinct in the wild in 1927, and a new founding population of 12 captive-bred individuals became the genetic basis for all extant bison (Węcek et al., 2017; Tokarska et al., 2011). As ecosystem engineers, they are thought to have a positive and significant impact on ecosystem remodelling. For example, the analysis done by Romero et al. (2015) highlighted that ecosystem engineers like European bison contribute to a 25% increase in species richness by creating and modifying habitats. They also play a role in shaping soil microbial communities, particularly through their dung, thereby contributing to soil nutrient cycling and energy flow within ecosystems (Lacher et al., 2019). Bison grazing can influence the composition and function of soil fauna communities, which are crucial for soil health and nutrient cycling (Ivanova et al., 2018). For example, research in the Kaluzhskie Zaseki State Nature Reserve, Russia, found that bison grazing led to differences in earthworm density and biomass across different habitats, with higher densities observed in certain transition zones between meadows and forests (Ivanova et al., 2018).

Given the population densities, associated infrastructure and consequent environmental pressures, one might question whether the UK is a suitable location for rewilding development (Loth & Newton, 2018; Ceaușu et al., 2015; Monbiot, 2015). However, this is precisely the reason that many advocates are pushing for rewilding in this country—in an effort to allow nature to flourish on this crowded island (Monbiot, 2013). Over the last decade we have seen extensive rewilding experimentation and engagement here. For example, the Knepp Castle Estate which located in West Sussex, UK, introduced a variety of large animals, including longhorn cattle, Exmoor ponies, red deer and wild boar in 2001, and stopped the use of pesticides and fertilisers, allowing wild plants to grow naturally (Knepp, n.d.; Tree, 2017). Another project is Vincent Wildlife Pine Marten Recovery Project in Wales. Between 2015 and 2021, 86 pine martens were reintroduced to the Forest of Dean and Wye Valley in Gloucestershire (Vincent Wildlife Trust, 2015).

Focusing on the Wilder Blean project, the Kent Wildlife Trust, in partnership with the Wildwood Trust, have embarked on a first for UK conservation: introducing European bison as natural woodland management in West Blean and Thornden Woods (Kent Wildlife Trust, n.d.). West Blean and Thornden Woods is a 781 hectare (1,930 acre) Site of Special Scientific Interest (SSSI) in north Canterbury, Kent (Natural England, n.d.). It is part of the Blean Woods Nature Reserve Review Site (a Grade I site), of which 490 hectares (1,200 acres) is a nature reserve (Ratcliffe, 2012), managed by the Kent Wildlife Trust. The forest is one of the largest areas of ancient woodland in the UK, with parts of the woodland more than 1,000 years old. In 2022, four adult European bison were introduced to West Blean and Thorndon Woods as part of the Wilder Blean project, and in the same year, one calf was born on the site, followed by another the year after. Each member of the herd has been specially selected to ensure a rich genetic pedigree (Kent Wildlife Trust, n.d.). The birth of the calves also marks a measurable advance forward in the project’s species recovery.

Supporters emphasise the potential of rewilding to create opportunities for ecosystem restoration while generating ecological and societal benefits. However, rewilding has also attracted criticism owing to uncertainty surrounding both its definition and its ecological outcomes, particularly over longer timescales (Perino et al., 2019; Prior & Ward, 2016; Hayward et al., 2019). A major challenge in evaluating rewilding interventions is that evidence has been derived predominantly from studies of vegetation and vertebrate communities, whereas belowground responses remain comparatively poorly understood. Soil microbial communities underpin key ecosystem functions, including nutrient cycling, decomposition and soil formation, and may respond rapidly to ecological change. Yet research on microbial diversity, biogeographic patterns and functional potential in rewilding systems has lagged considerably behind studies of macroflora and fauna (Perino et al., 2019). Consequently, important uncertainties remain regarding how rewilding influences ecosystem processes at the microbial level and how changes in microbial functional genes translate into broader shifts in community structure and ecosystem functioning under changing environmental conditions (Louca et al., 2016; Breitkreuz et al., 2021). Addressing these knowledge gaps is critical for developing a more mechanistic understanding of rewilding outcomes and for evaluating whether rewilding interventions deliver the ecosystem changes that they seek to promote.

Accordingly, this study aimed not only to determine whether the introduction of European bison was associated with early changes in the taxonomic composition and functional potential of forest soil communities in West Blean and Thornden Woods, but also to fill in the gap in research on microbial diversity, biogeographic patterns and functional potential in rewilding systems. Using Oxford Nanopore Technologies (ONT) shotgun metagenomic sequencing, we compared soil communities before and two years after bison introduction, incorporating both repeated temporal sampling and contemporaneous comparisons across different grazing and management regimes. ONT sequencing generates long reads, which substantially improves metagenomic assembly continuity, reducing the fragmentation commonly associated with short-read sequencing platforms (Latorre-Pérez et al., 2020). Consequently, this approach may offer greater sensitivity for detecting subtle ecological changes over relatively short timescales, making it particularly well suited for investigating the early functional responses of soil microbial communities to rewilding. We hypothesised that the introduction of bison would be associated with detectable shifts in soil community composition and functional potential in the Bison Area relative to both its pre-introduction state and the other management areas. We further hypothesised that, during this early stage of rewilding, changes in functional composition would be more pronounced than changes in taxonomic alpha diversity, reflecting a potentially earlier response of microbial functional potential to ecological disturbance.

## 2. Material and methods

### 2.1 Sampling

The area of West Blean and Thornden Woods was divided into three areas, the Control Area, Grazer Area and Bison Area (Figure 1). A total of 56 soil samples were selected for the metagenomic analysis out of a total of 140 collected in 2022 and 2024 from three areas in West Blean and Thornden Woods before the European bison were released in 2022 (28 samples) and 2024 (28 samples). The location and number of samples were dictated by the even distribution across the woodland in 200 meter distance from each other. From each year’s batch, 10 samples were taken from the Control area, 10 from the Grazer area, and 8 from the Bison area (Figure 1). The soil samples were collected using a core borer at a depth of 30 cm at each sampling site to minimise environmental perturbations to the community structure. The core borer was carefully wiped with 80% ethanol after each sample was taken to limit cross-contamination between samples. All 2022 soil samples and 2024 samples from the bison area were collected by Kent Wildlife Trust team, other 2024 samples were collected by the Natural History Museum team. After DNA extraction, the samples were transferred to a −20°C freezer until further molecular analyses.

**Figure 1.**
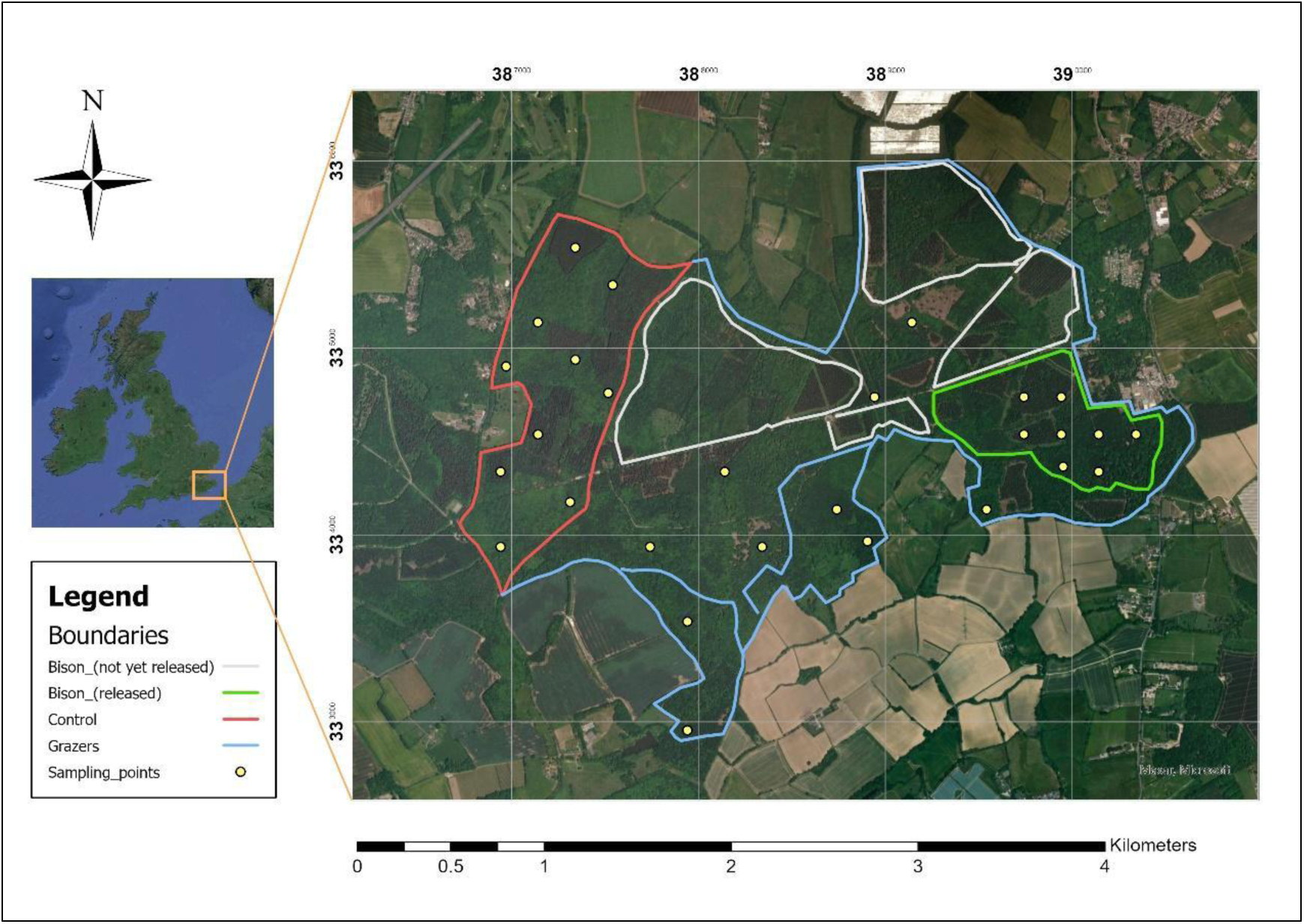
Location of the study area and distribution of soil sampling points in West Blean and Thornden Woods, Kent, UK. The map of the areas was generated using ArcGIS pro 3.0.0. The westernmost Control Area covers approximately 100ha, it remains under human maintenance and serves as a control site. Within the blue area is the Grazer Combined Area, which covers 260ha. Domestic grazers: 5 Longhorn cattle, 6 Exmoor ponies and 3 Iron Age pigs were released in this area. They were already present in 2022. The green area is an area designated for European Bison and covers approximately 205ha. Konik ponies were present in this area, while the first four European bison replaced them in 2022.

### 2.2 DNA extraction, preprocessing and ONT sequencing

DNA was extracted from the selected soil samples using the DNeasy PowerSoil Pro Kit (Qiagen). The extracted DNA samples were then additionally cleaned and purified using the DNeasy PowerClean Pro Cleanup Kit (Qiagen). The purity of the extracted DNA was determined using NanoDrop spectrophotometer (Thermo Fisher Scientific). Qubit Fluorometer (Thermo Fisher Scientific) was used to accurately measure the concentration of the extracted DNA. 1% agarose gel electrophoresis was used to estimate the DNA fragment sizes. DNA was extracted in triplicate from each soil sample to account for variation in extraction efficiency and technical error, yielding a total of 168 DNA extracts for sequencing. Additionally, 12 negative controls containing molecular grade water used for the extractions of DNA samples were also prepared alongside technical replicates, 6 for technical replicates from 2022 and 6 for year 2024. The purified DNA samples were used for library preparation using Oxford Nanopore Technologies Rapid barcoding 96 V14 kit (SQK-RBK114.96) following Rapid Sequencing DNA V14 protocol dedicated to this kit. Libraries were pooled by sampling year (2022 and 2024) and sequenced on separate PromethION flow cells (Oxford Nanopore Technologies).

### 2.3 Taxonomic classification

The sequencing reads obtained from the three sets of replicates for each sampling point were first concatenated into a single FASTQ file. Taxonomic classification and preprocessing of the shotgun metagenomic data were performed using the nf-core/taxprofiler pipeline v1.1.8 (Stamouli et al., 2023). Read quality was assessed using FastQC, adapter sequences were removed using Porechop, and low-quality reads were filtered using NanoFilt with a minimum quality score threshold of Q10. Taxonomic classification was subsequently performed using Centrifuge. A local Centrifuge database was constructed from the NCBI nucleotide non-redundant (nt) database (last updated on 16 May 2024) using the centrifuge-build utility. This database was subsequently used for taxonomic classification of the quality-filtered sequencing reads.

### 2.4 Normalization and relative abundance analysis

Transcripts Per Million (TPM) normalisation and relative abundance calculations were performed using a custom Python script (SI_1; SI_2). TPM normalisation is a common method in RNA-Seq data analysis and is also often used in metagenomic studies. Read counts were extracted from Centrifuge reports and normalised to account for differences in genome size and sequencing depth across samples. Relative abundances were subsequently calculated from the TPM-normalised read counts (SI_5).

The 56 samples were divided into six groups according to the year and location of collection: 2022 Control Area, 2022 Grazer Area, 2022 Bison Area, 2024 Control Area, 2024 Grazer Area, and 2024 Bison Area. The relative abundance matrices of the six sets of samples were analysed for alpha and beta diversity using the vegan package 2.6.6.1 in R version 4.3.3, and the taxonomic composition was visualized at the phylum level. Simpson index was used for alpha diversity assessment. Pairwise comparisons were performed using paired Wilcoxon signed-rank tests, whereas comparisons among three groups were performed using Kruskal–Wallis tests. Bray–Curtis dissimilarities were calculated to quantify differences in community composition, and PCoA was used to visualize the resulting distance matrix. ANOSIM was used to test whether community composition differed significantly among groups.

In addition, the relative abundance matrix was filtered to retain the seven most abundant nitrogen-fixing bacteria, eleven photosynthesising microbes and one mycorrhizal fungi at genus level, and performed non-parametric tests on the abundance of these genera between groups to quantify the effect of the presence of European bison on the abundance of these microorganisms. The relative abundance matrices were also screened to obtain all plant species. Simpson’s indices of vegetation diversity under the different groupings were calculated separately, and performed non-parametric tests to quantify the effect of the presence of European bison on plant diversity. Screening of plant species was done by matching our relative abundance matrix with the plant species list backbone file published by World Flora Online (WFO, 2023) using its corresponding WorldFlora R package 1.14.4 (Kindt, 2020). P-values of all tests were corrected using the Benjamini-Hochberg method.

### 2.5 Genome assembly and KEGG pathway annotation

The shotgun metagenomic sequencing data were assembled using MEGAHIT v1.2.9 using k-mer values ranging from 35 to 255. QUAST v5.2.0 was used to evaluate the assembly results. Gene prediction and KEGG annotation were performed using eggNOG-mapper v2.1.12. The R package vegan was used to calculate Bray–Curtis dissimilarities based on KEGG pathway abundance profiles to assess differences in pathway composition across different spatial and temporal scales. Principal coordinates analysis (PCoA) was performed to visualize the overall variation in pathway composition among groups, while permutational multivariate analysis of variance (PERMANOVA) was used to statistically assess differences in pathway composition between groups. Differentially abundant KEGG pathways were identified using the R package DESeq2 (version 1.40.2), followed by KEGG pathway enrichment analysis. The enriched pathways were visualized using ggplot2.

## 3. Results

### 3.1 General characteristics of the soil microbial communities

The sequencing runs yielded 25.7 million raw reads comprising an estimated 41.17 Gb of sequence data for the 2022 samples and 27.29 million raw reads representing 40.77 Gb of sequence data for the 2024 samples. Read lengths were predominantly between 600 bp to 1200 bp. The number of total reads per barcode varied from 3.25 Mb to 845.75 Mb, highlighting the importance of normalization. Across all samples, 8,689,711 reads were successfully classified. A total of 108 phylum-level taxa were identified across the 56 samples. No taxa were detected in the negative controls, indicating an absence of detectable contamination during sample processing. Figure 2 shows the relative abundance of taxa at the phylum level in each sample, with only the ten most abundant phyla displayed. Overall, community structure was broadly similar across all groups. Proteobacteria and Actinobacteria dominated all 56 samples, together accounting for more than 75% of the total relative abundance. These were followed by Chordata, with an abundance ranging from 3% to 7%. The remaining phyla each contributed relatively small proportions of the overall community composition.

**Figure 2.**
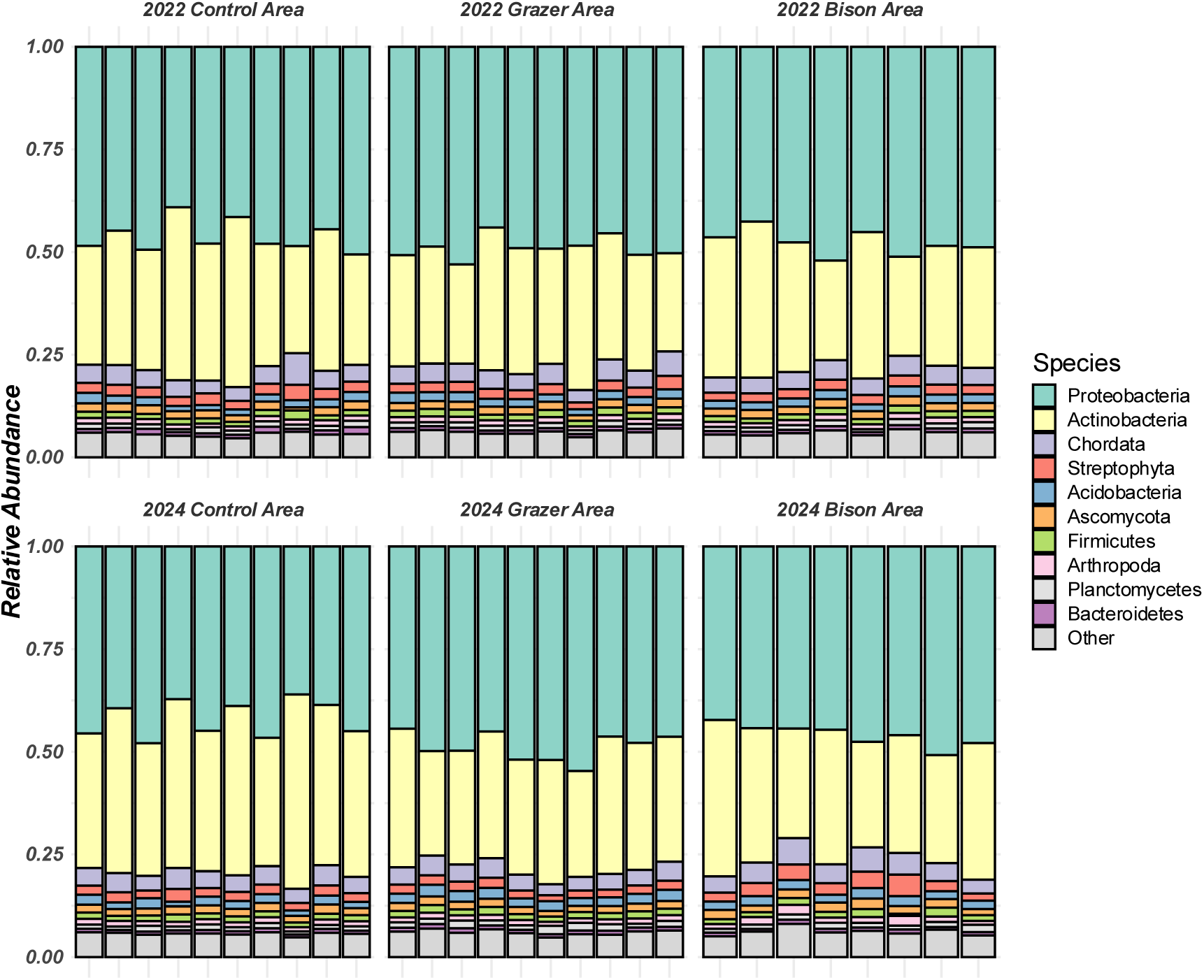
Relative abundances of the ten most abundant phyla across soil samples collected from the Control, Grazer and Bison areas in 2022 and 2024. Each stacked bar represents one soil sample.

### 3.2 Alpha diversity

Figure 3 shows the temporal and spatial variation in Simpson’s diversity index across the different study areas. The median Simpson’s index exceeded 0.9 in all areas, indicating consistently high levels of biodiversity. In both the Control and Bison areas, median Simpson’s index values were slightly lower in 2024 than in 2022 Bison Area. Among the 2022 sites, the Bison Area exhibited a lower Simpson’s index than the Control Area but a higher index than the Grazer Area. In contrast, the Bison Area had the highest median Simpson’s index of the three areas in 2024. The Grazer Area consistently showed the lowest alpha diversity index of the three areas in both years, despite grazers having been introduced before 2022. However, non-parametric tests for the five groups of comparisons showed non-significant differences (p_adjusted>0.05) for all comparisons.

**Figure 3.**
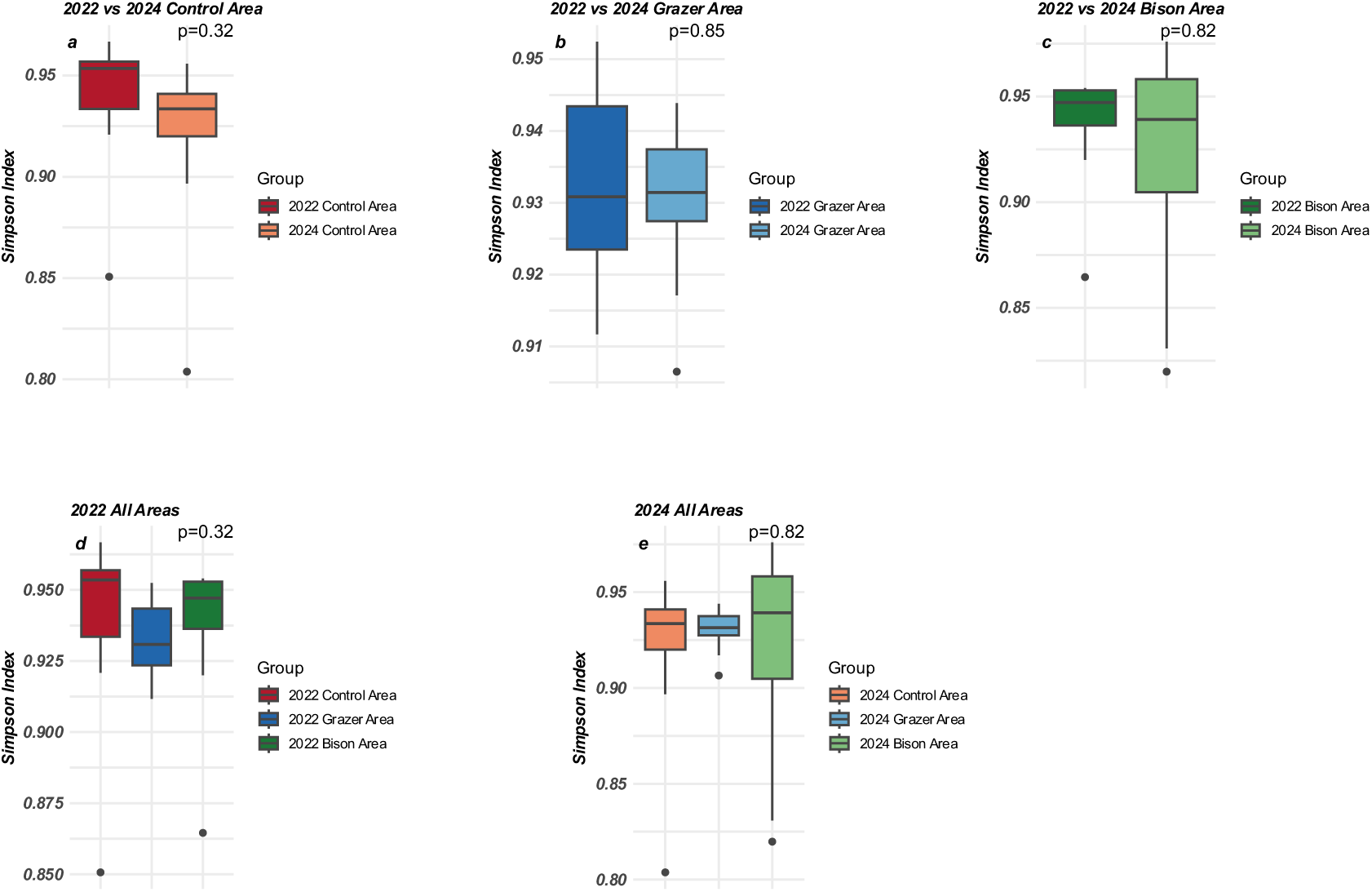
Temporal and spatial comparisons of taxonomic alpha diversity, measured using Simpson’s diversity index, among the Control, Grazer and Bison areas. Panels show comparisons between 2022 and 2024 in the (a) Control Area, (b) Grazer Area and (c) Bison Area, and comparisons among the Control, Grazer and Bison areas in (d) 2022 and (e) 2024.

### 3.3 Beta diversity

Figure 4 illustrates differences in community composition among the study areas. Samples from the 2022 Bison Area formed a relatively tight cluster, indicating low within-group variation, whereas samples from the 2024 Bison Area were more dispersed, reflecting greater within-group variation. The 2024 Bison Area was also clearly separated from the 2022 Bison Area on both PCoA1 and PCoA2. The taxonomic community composition of the 2024 Bison Area differed significantly from that of the 2022 Bison Area (R=0.37, p<0.05). ANOSIM detected a significant overall difference among the three 2024 treatment areas (R = 0.20, P < 0.05), although no significant pairwise differences remained after correction for multiple testing.

**Figure 4.**
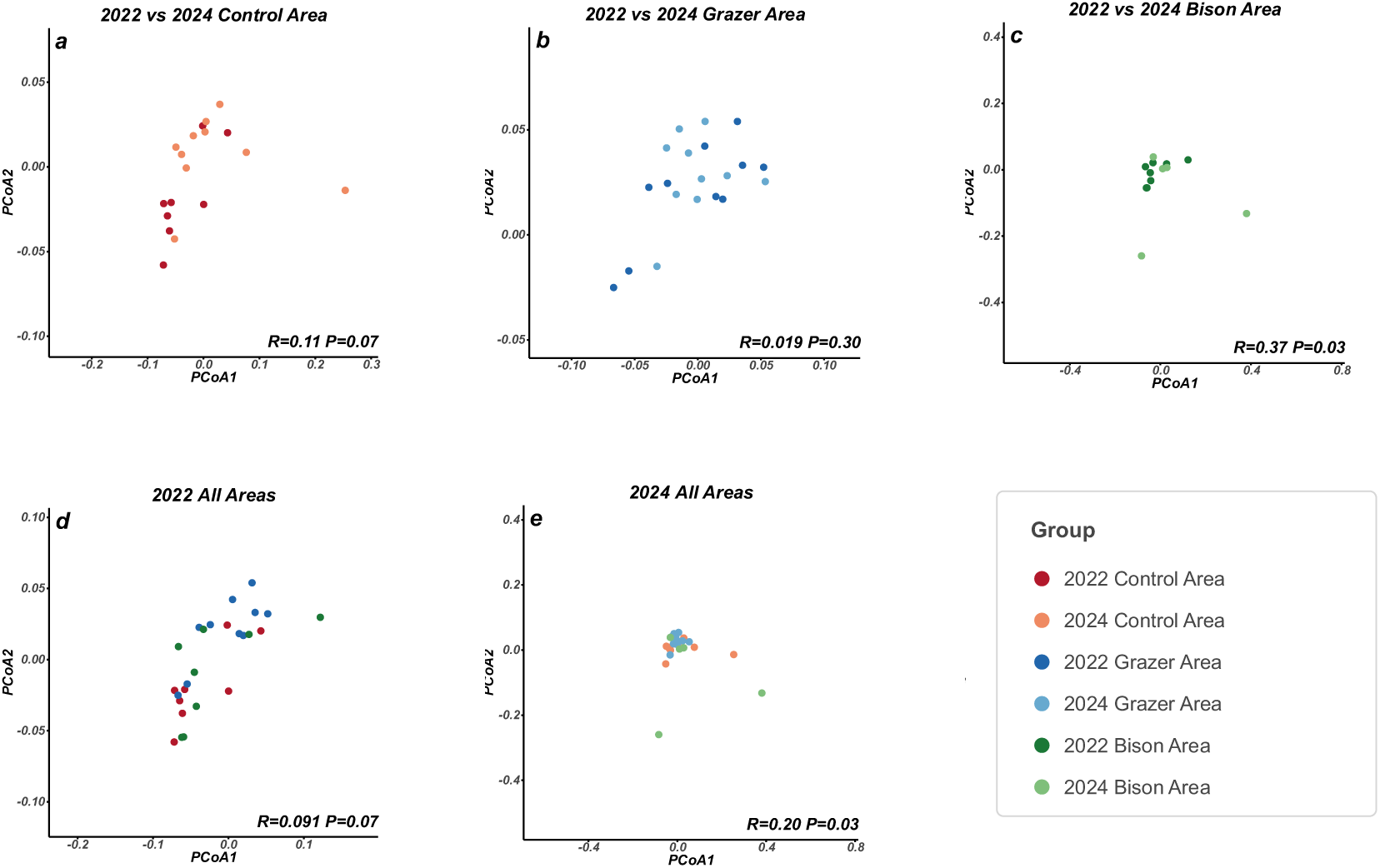
Principal coordinates analysis of taxonomic community composition based on Bray– Curtis dissimilarities among soil samples from the Control, Grazer and Bison areas. Panels show comparisons between 2022 and 2024 in the (a) Control Area, (b) Grazer Area and (c) Bison Area, and comparisons among the Control, Grazer and Bison areas in (d) 2022 and (e) 2024.

The greater dispersion observed in the 2024 Bison Area was consistent with the pattern identified for Simpson’s diversity index (Figure 3), where samples from this area also exhibited greater variability than those from other groups. Together these results suggest increased heterogeneity in community composition within the Bison Area between 2022 and 2024.

### 3.4 Spatial and temporal difference of rhizobia and photosynthetic microorganisms

Spatial and temporal variations in the relative abundance of the top seven genera of rhizobia and the top eleven of photosynthetic microorganisms were compared. Figure 5 illustrates the variation of the seven rhizobial genera that contain known symbiotic nitrogen-fixing representatives, namely *Azorhizobium*, *Mesorhizobium*, *Neorhizobium*, *Rhizobium*, *Bradyrhizobium*, *Pararhizobium* and *Sinorhizobium*. Non-parametric tests showed that the relative abundance of *Azorhizobium* was significantly lower (p<0.05) in the 2024 Bison Area compared to the same area in 2022 (Figure 5c). Among the three areas in 2024, the relative abundance of Azorhizobium in the Bison Area was significantly lower (p<0.05) than the other two areas (Figure 5e). Although the remaining genera did not differ significantly between groups, *Azorhizobium*, *Mesorhizobium*, *Neorhizobium*, *Rhizobium*, and *Sinorhizobium* all exhibited lower median relative abundances in the 2024 Bison Area than in the corresponding 2022 samples (Figure 5c). These five genera also exhibited their lowest median relative abundances in the 2024 Bison Area compared with the other treatment areas (Figure 5e). In contrast, little variation in the relative abundance of the seven rhizobial genera was observed between the 2022 and 2024 Control areas (Figure 5a), between the 2022 and 2024 Grazer areas (Figure 5b), or among the three treatment areas in 2022 (Figure 5d).

**Figure 5.**
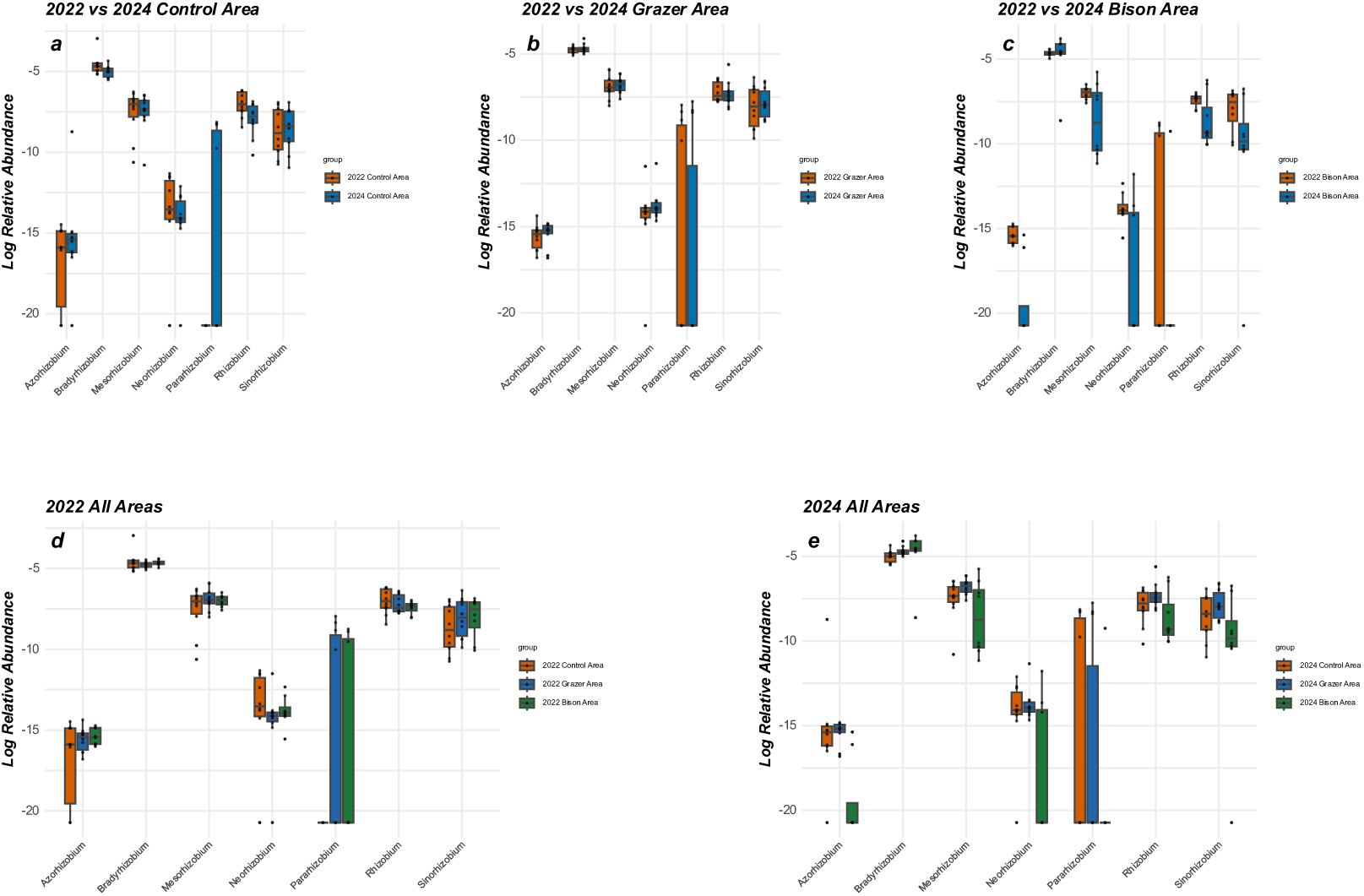
Temporal and spatial variation in the relative abundances of seven bacterial genera containing known symbiotic nitrogen-fixing representatives. Panels show comparisons between 2022 and 2024 in the (a) Control Area, (b) Grazer Area and (c) Bison Area, and comparisons among the Control, Grazer and Bison areas in (d) 2022 and (e) 2024.

Figure 6 illustrates the spatial and temporal variation in the relative abundance of the top eleven genera of microorganisms containing known photosynthetic representatives, namely *Anabaena*, *Chlorobium*, *Chloroflexus*, *Nostoc*, *Prochlorococcus*, *Pseudanabaena*, *Rhodobacter*, *Rhodopseudomonas*, *Rhodospirillum*, *Synechococcus* and *Oscillatoria*. Non-parametric tests showed that none of the photosynthetic microbial genera were significantly different in relative abundance at the statistical level. Nevertheless, from Figure 6 we can still see that in the 2024 Bison Area (Figure 6c), except for *Oscillatoria,* the remaining ten genera of photosynthetic microbial genera (*Anabaena, Chlorobium, Chloroflexus, Nostoc. Prochlorococcus, Pseudanabaena, Rhodobacter, Rhodopseudomonas, Rhodospirillum, Synechococcus*) all exhibited lower relative abundances than in the 2022 Bison Area. This pattern was not found in the inter-annual comparisons of both the Control Area (Figure 6a) or the Grazer Area (Figure 6b). In the comparison of the three areas of 2024 (Figure 6e), although these differences were not statistically significant, nine of the eleven genera had the lowest median relative abundance in the 2024 Bison Area (*Anabaena, Chlorobium, Chloroflexus, Prochlorococcus, Pseudanabaena, Rhodobacter Rhodopseudomonas, Rhodospirillum, Synechococcus*). In contrast, this pattern did not occur in the 2022 comparison among the three areas (Figure 6d). This variation is similar to the changes in rhizobial genera described in the previous section (Figure 5).

**Figure 6.**
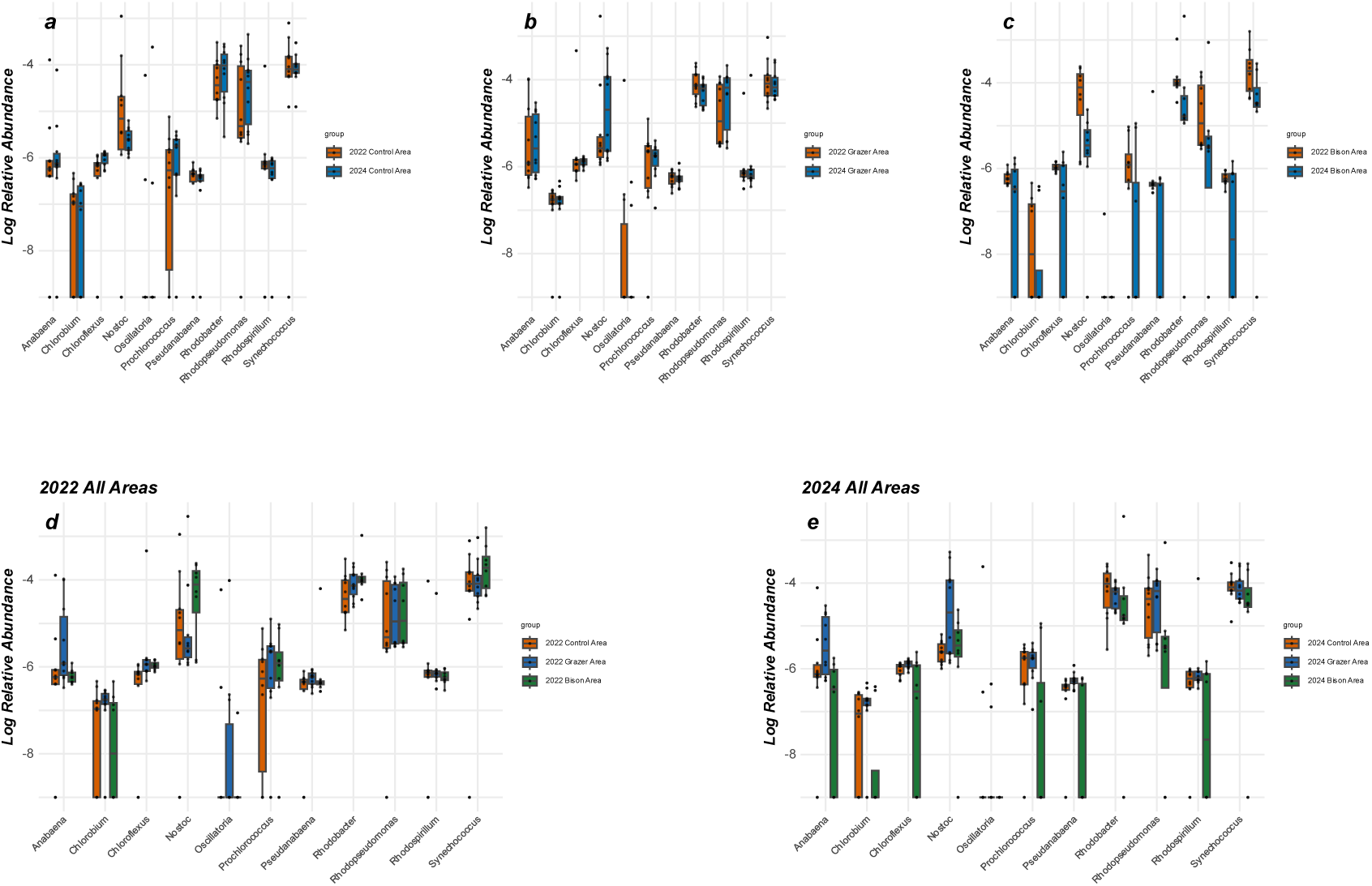
Temporal and spatial variation in the relative abundances of eleven microbial genera containing known photosynthetic representatives. Panels show comparisons between 2022 and 2024 in the (a) Control Area, (b) Grazer Area and (c) Bison Area, and comparisons among the Control, Grazer and Bison areas in (d) 2022 and (e) 2024.

### 3.5 Spatial and temporal difference of vegetation and mycorrhizal fungi

A total of 3652 plant taxa were detected from the relative abundance matrix of species obtained from metagenomic sequencing. The calculation of Simpson’s index of metagenomically detected plant taxon diversity under different groupings is shown in Figure 7. Non-parametric tests showed that no significant difference was observed for all five groups of comparison (p>0.05). Figure 8 shows the spatial and temporal variations in the relative abundance of the mycorrhizal genus *Rhizophagus*. Non-parametric tests showed that the relative abundance of Rhizophagus was significantly higher in the 2024 Control Area than in the 2022 Control Area (p<0.05 [Figure 8a]), and significantly lower in the 2024 Grazer Area and the 2024 Bison Area than in the 2024 Control Area (p<0.05 [Figure 8e]).

**Figure 7.**
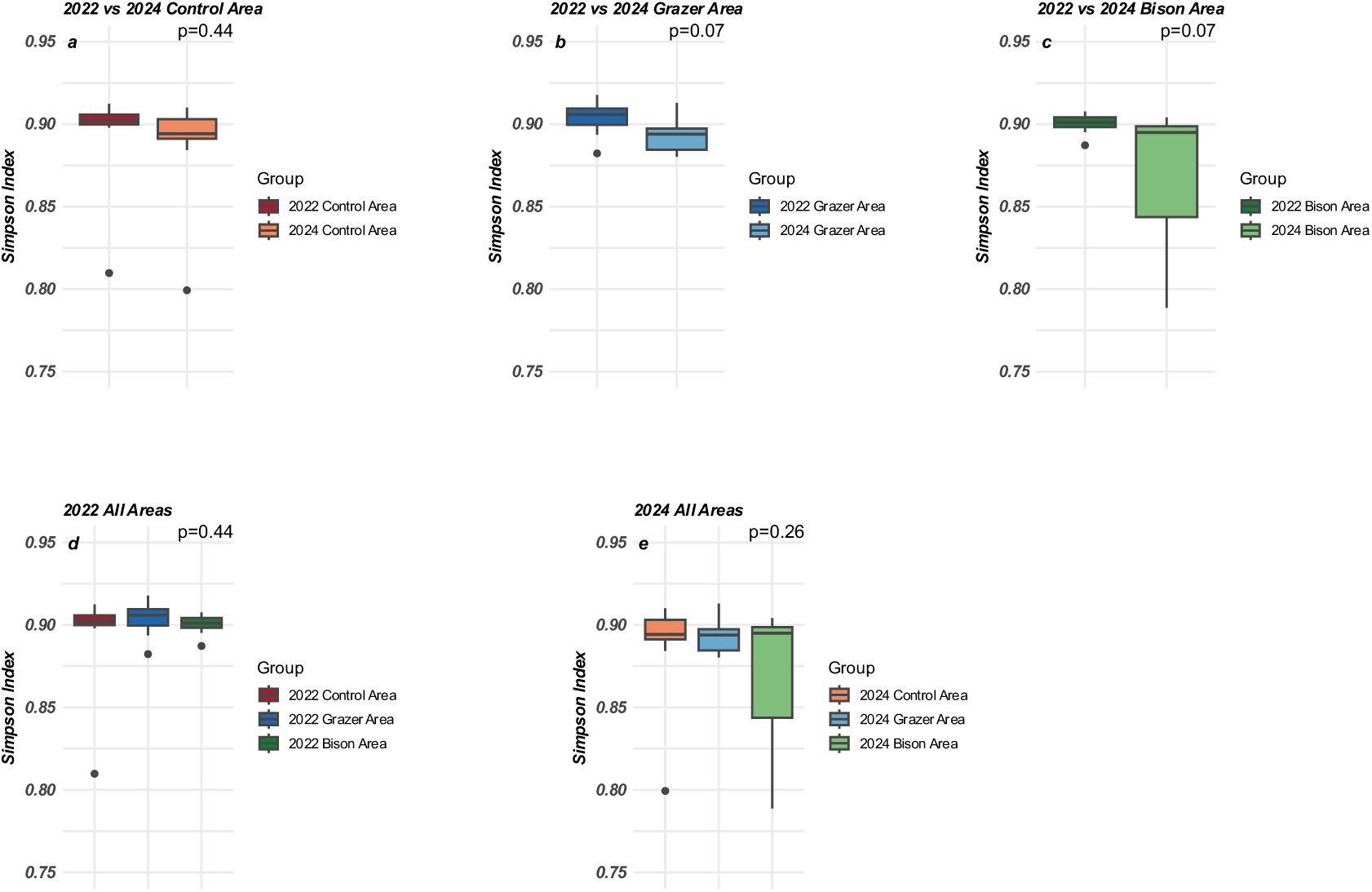
Temporal and spatial comparisons of the diversity of metagenomically detected plant taxa, measured using Simpson’s diversity index. Panels show comparisons between 2022 and 2024 in the (a) Control Area, (b) Grazer Area and (c) Bison Area, and comparisons among the Control, Grazer and Bison areas in (d) 2022 and (e) 2024.

**Figure 8.**
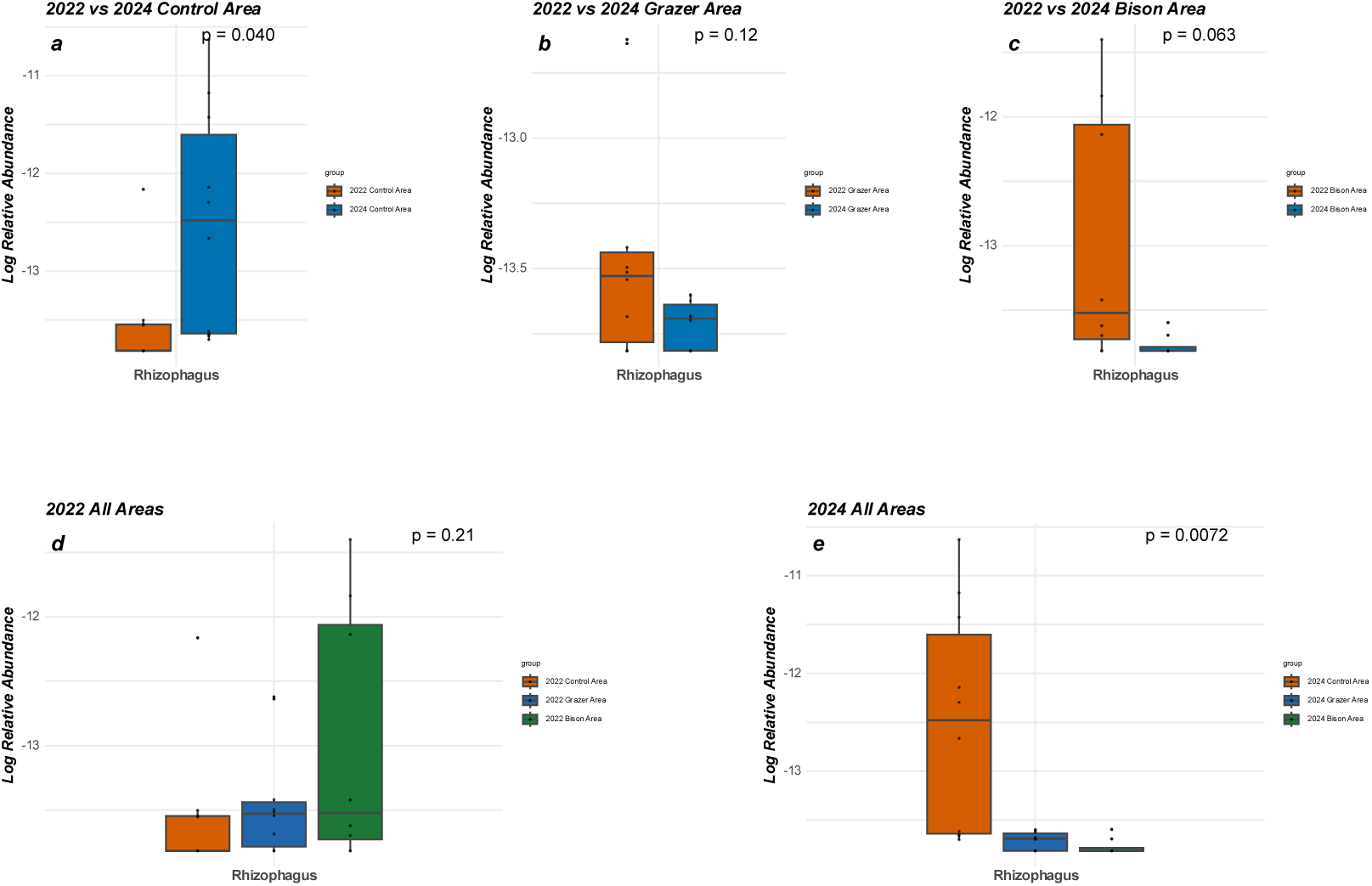
Temporal and spatial variation in the relative abundance of the arbuscular mycorrhizal fungal genus *Rhizophagus*. Panels show comparisons between 2022 and 2024 in the (a) Control Area, (b) Grazer Area and (c) Bison Area, and comparisons among the Control, Grazer and Bison areas in (d) 2022 and (e) 2024.

### 3.6 Functional beta diversity of KEGG pathways

PCoA based on Bray–Curtis dissimilarities of KEGG pathway abundance profiles revealed distinct separation in functional composition between the 2024 Bison area and the other groups (Figure 9). PERMANOVA further confirmed significant differences in KEGG pathway composition between the 2024 Bison area and the 2022 Bison area (R² = 0.35, F = 7.53, P = 0.001), the 2024 Control area (R² = 0.26, F = 5.63, P = 0.008), and the 2024 Grazer area (R² = 0.33, F = 7.75, P = 0.002) (Table 1). These results indicate that the functional composition of soil communities in the 2024 Bison area differed significantly from those in the corresponding comparison groups. In all three pairwise comparisons, the 2024 Bison area showed distinct functional profiles from the corresponding comparison groups.

**Figure 9.**
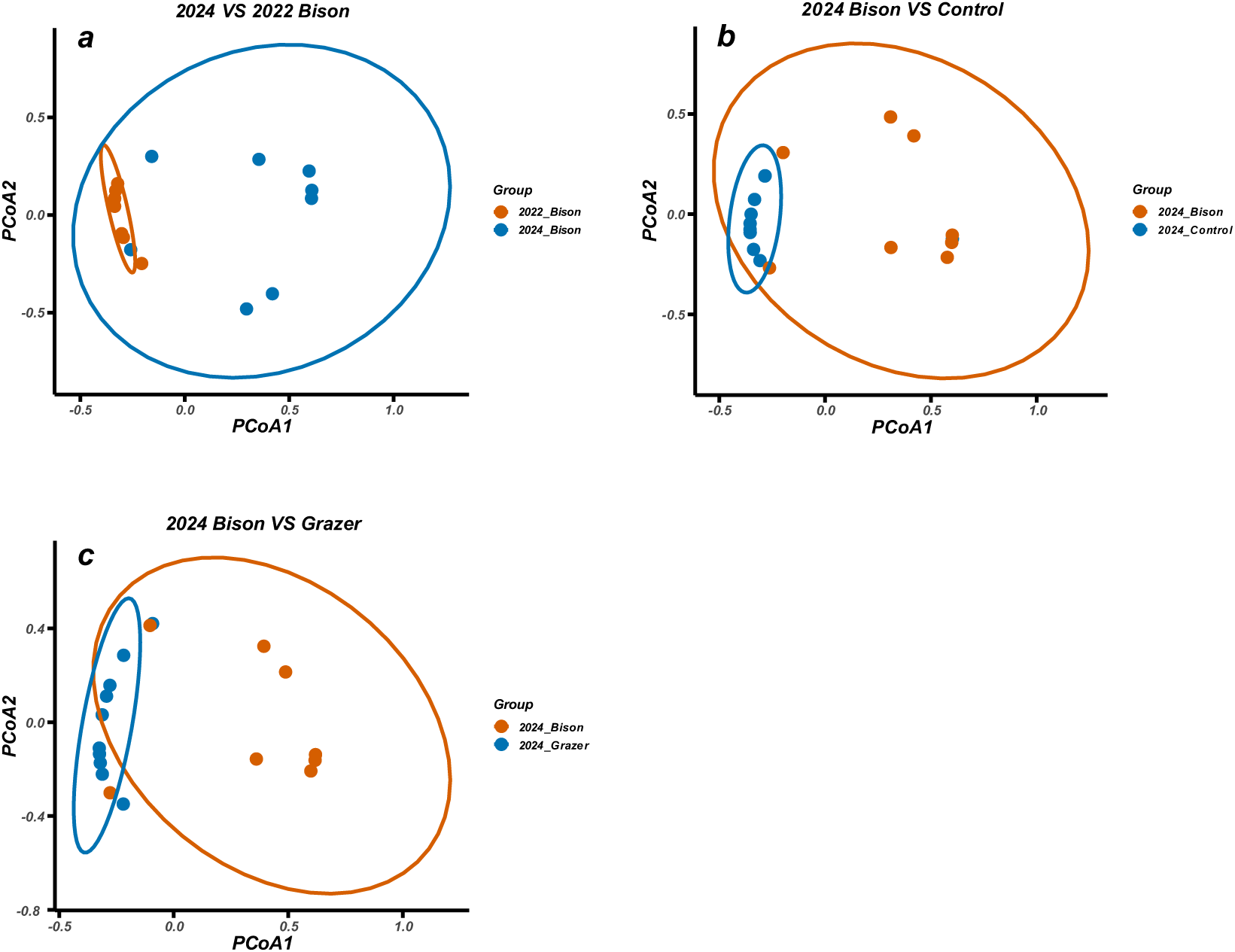
Principal coordinates analysis of functional beta diversity based on Bray–Curtis dissimilarities of KEGG pathway abundance profiles. Panels compare the 2024 Bison Area with (a) the 2022 Bison Area, (b) the 2024 Control Area and (c) the 2024 Grazer Area.

**Table 1.** PERMANOVA results for pairwise differences in functional composition based on KEGG pathway abundance profiles.

|  | **R<sup>2</sup>** | **F value** | **P value** |
| --- | --- | --- | --- |
| **2024 Bison VS 2022 Bison** | 0.35 | 7.53 | 0.001 |
| **2024 Bison VS 2024 Control** | 0.26 | 5.63 | 0.008 |
| **2024 Bison VS 2024 Grazer** | 0.33 | 7.75 | 0.002 |

### 3.7 KEGG pathway enrichment analysis

KEGG pathway enrichment analysis revealed both shared and comparison-specific functional shifts among the groups (Figure 10). Carbon metabolism and amino acid biosynthesis were consistently enriched in the 2024 Bison area compared with the 2022 Bison area, the 2024 Control area, and the 2024 Grazer area. In addition, fatty acid metabolism was enriched in the comparison between the 2024 and 2022 Bison areas, whereas pyruvate metabolism was enriched between the 2024 Bison and 2024 Control areas. Quorum sensing was uniquely enriched in the comparison between the 2024 Bison and 2024 Grazer areas.

**Figure 10.**
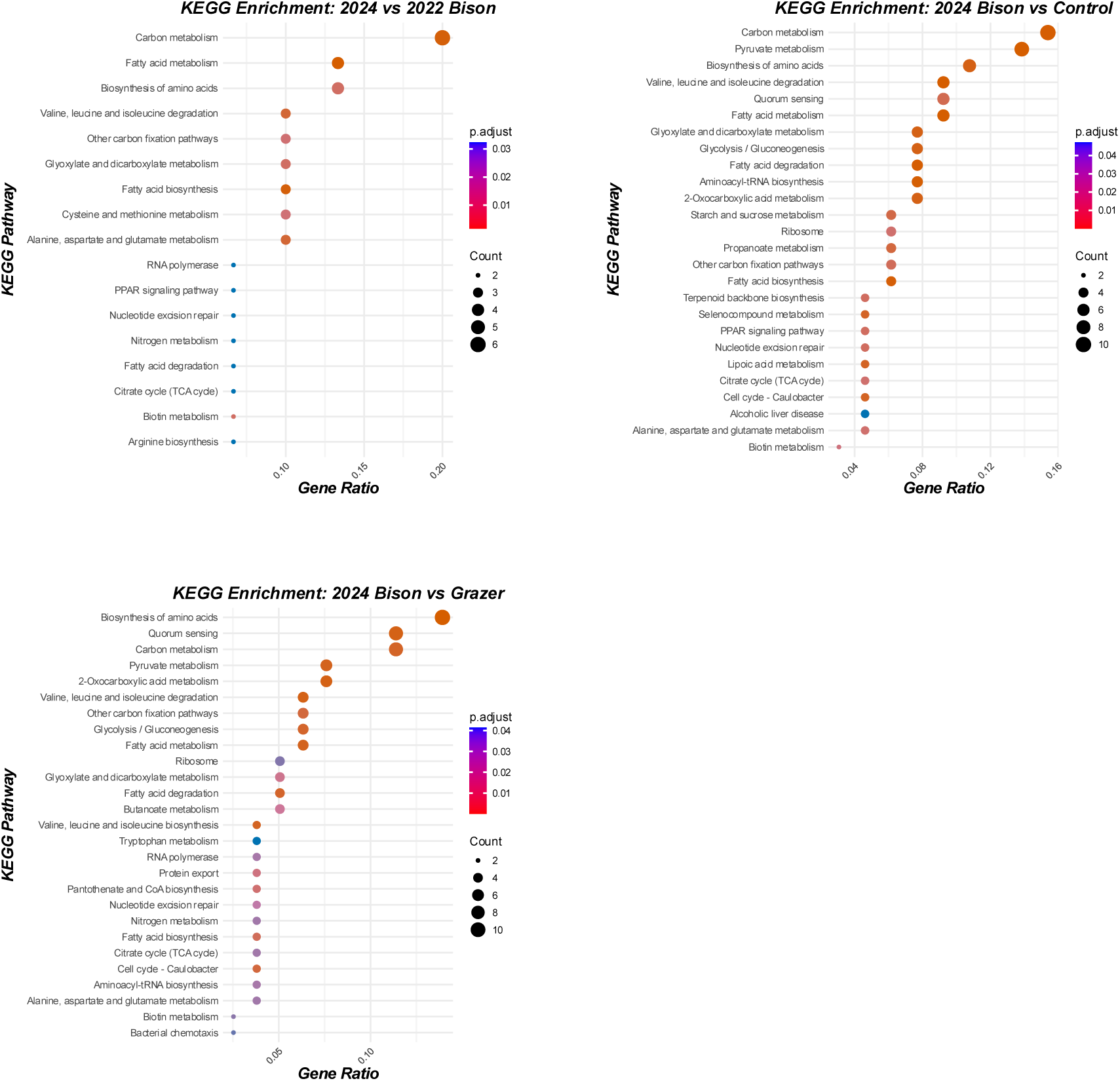
KEGG pathway enrichment analysis of functional differences involving the 2024 Bison Area. Dot plots show enriched KEGG pathways identified following DESeq2-based differential-abundance analysis for comparisons between (a) the 2024 and 2022 Bison areas, (b) the 2024 Bison and 2024 Control areas, and (c) the 2024 Bison and 2024 Grazer areas.

## 4. Discussion

### 4.1 Effect of the presence of European bison on the taxonomic diversity of soil biomes

Two years after the reintroduction of European bison, soil microbial community composition in the Bison Area differed significantly from its pre-reintroduction state in 2022, although taxonomic alpha diversity remained largely unchanged. This indicates that the observed temporal response was characterised primarily by shifts in community composition rather than by detectable changes in overall diversity. However, no significant differences in community composition were detected among the Bison, Grazer and Control areas in 2024. Therefore, these results do not provide strong evidence that the observed changes were specifically attributable to the reintroduction of bison, rather than to other environmental factors. Overall, these findings do not support our hypothesis proposed above, as there is currently not enough evidence to conclude that the reintroduction of a keystone species increased the taxonomic diversity of the soil biotic community. Although some previous studies had reported the opposite conclusion, for example, in tallgrass prairies, the introduction of American bison dung has been shown to increase microbial diversity in the soil (Hawkins & Zeglin, 2022). This suggests that bison act as vectors for microbial dispersal, contributing to a more homogenised and diverse microbial community across different land management practices. The research by Ivanova et al. (2018) examines how the free grazing of European bison affects vegetation and soil fauna, including earthworm populations, contributing to increased soil biodiversity. However, they also noted that such significant changes would take 10-15 years (Ivanova et al., 2018). Similarly, in our study, the two-year period may not be sufficient to significantly affect soil alpha diversity, especially in stable soil environments. Even if beta diversity changes rapidly due to the disturbance of a new environment, alpha diversity may take longer to show significant changes (Flohre et al., 2011). In addition to this, if soil resources (e.g., nutrients, moisture) in the study area were originally adequate and homogeneous, the microbial community may have maintained its original high level of alpha diversity after the introduction of bison (Prober et al., 2015). This is supported by the fact that the median of Simpon’s index of all six groups of samples in this study was greater than 0.9, which represents the high levels of alpha diversity.

Although there was no increase in the number of species, the change in beta diversity suggests that the soils with the presence of bison did differ from other soils in terms of community structure. Such differences may be manifested in the substitution of species types or changes in the relative abundance of the same species, i.e., the species composition of the bison region was more heterogeneous. These phenomena may be due to local environmental perturbations caused by the introduction of European bison, such as increased soil heterogeneity through trampling and herbivory. The migratory behaviour of bison and the distribution of faeces may also lead to increased heterogeneity of soil microbial communities (Augustine & Frank, 2001; Jaroszewicz, 2013). This heterogeneity includes both differences within the Bison Area and differences between the Bison Area and other Areas. Although these changes may not be sufficient to significantly alter overall species richness (i.e. alpha diversity), they can lead to an increase in differences in species composition between sites, which in turn can lead to an increase in beta diversity (Barber et al., 2023). This interpretation is consistent with the greater dispersion of samples of the 2024 Bison Area samples shown in the Beta diversity analysis. Fukami & Nakajima (2013) pointed out that these behaviours of bison increased the diversity of soil conditions in localised areas, but the overall species diversity of soil microbes (alpha diversity) may not have been significantly affected. Similarly, another study showed that in a marine benthic community in a rocky pond in the St. Lawrence Estuary, Canada, low intensity trophic disturbance significantly increased the community’s beta diversity, although it had little effect on alpha diversity (Séguin et al., 2013). This study also suggests that beta diversity may be more sensitive to environmental changes and therefore better indicative of minor biodiversity changes (Séguin et al., 2013). In addition to this, when the spatial scale increased from local to regional, it was found that there was a switch from beta diversity dominance to alpha diversity dominance (Gering & Crist, 2002). That is, as spatial scale increases, the contribution of beta diversity to total diversity increases, and species differences between sites within the region are dominating total species richness.

However, soil alpha diversity is influenced by a variety of factors such as soil physicochemical properties (e.g. water content, pH, total carbon, total nitrogen, carbon to nitrogen ratio), climatic conditions (e.g. mean annual temperature, precipitation), etc. (Wang et al., 2015; Zhao et al., 2022). Therefore, in the future, more comprehensive integrated measurements of soil variables in the study area are likely to be able to provide a more convincing explanation for the current state of soil taxonomic diversity.

### 4.2 Changes in soil functional diversity following bison rewilding

Enrichment of amino acid synthesis pathways appeared in the post-rewilding Bison Area in comparison to the Control Area, the Grazer Area and the pre-rewilding Bison Area. Since amino acid synthesis is directly dependent on available nitrogen sources in the soil (Zhang et al., 2015; Jämtgård et al., 2010), this result is consistent with the possibility that nitrogen availability or nitrogen inputs increased following bison introduction, although direct measurements of soil nitrogen are required to confirm this interpretation. This hypothesis is also supported by the decrease in nitrogen-fixing bacteria observed in the soils of the Bison Area. Due to the increase in soil organic nitrogen sources, the dependence of plants on symbiotic nitrogen-fixing bacteria for nitrogen fixation from air was reduced. The dominance of Proteobacteria may be consistent with relatively nutrient-rich conditions in the soil, as they are ‘eutrophic’ microorganisms that are able to rapidly utilise the rich carbon and nitrogen sources in the soil for reproduction (Spain et al., 2009). Bison manure and plant disturbance are important sources of nitrogen input. Bison activity can introduce large amounts of organic and inorganic nitrogen into the soil through dung, urine, and vegetation disturbance, thereby increasing the pool of available nitrogen in the soil (Hawkins & Zeglin, 2022). This may also have a positive impact on the overall nutrient cycling of the soil ecosystem.

The enrichment of amino acid biosynthesis relative to the Grazer Area may reflect that, although other herbivores are present in the Grazer Area, their activity patterns and nitrogen input modes (e.g., excretion modes) may have been different from those of bison, resulting in a more dispersed distribution of nitrogen in the soil (Pruszenski & Hernández, 2020), which may have reduced localised nitrogen concentration, leading to less significant enrichment of amino acid synthesis pathways than in the Bison Area. Second, the significantly larger size of the Grazer Area than the Bison Area may have resulted in more spatially diffuse nitrogen inputs. Grazing intensity may also have differed between the areas: the 205-ha Bison Area supported six bison during 2024, equivalent to approximately 0.029 bison ha⁻¹, whereas the 260-ha Grazer Area supported a temporally varying mixture of pigs, Exmoor ponies and Longhorn cattle. Although these raw densities are not directly comparable across species, differences in effective stocking rate and excreta distribution may have contributed to the observed functional differences.

The enrichment of carbon metabolic pathways in the three comparative groups reflects the activity of microorganisms in utilising and transforming carbon resources. Enriched carbon metabolic pathways can include both carbon fixation and carbon decomposition processes. However, the enrichment of the pyruvate metabolic pathway when 2024 Bison Area was compared with 2024 Control Area and 2024 Grazer Area may indicate increased potential for microbial carbon turnover. This is due to the fact that pyruvate metabolism is an important intermediate step in the glycolysis and tricarboxylic acid cycles, which typically involves further oxidation of carbon and release of energy (Marín-García, 2013; Ha & Bhagavan, 2023). The enrichment of fatty acid metabolic pathways when comparing the 2024 Bison Area to the 2022 Bison area also typically indicates that soil microorganisms are actively processing carbon resources. This result differs from most previous studies, for example, multiple studies highlighted how the reintroduction of bison has led to a significant increase in plant diversity, which in turn contributes to greater soil carbon storage (Ratajczak et al., 2022; Chen et al., 2018; Lange et al., 2015). One possible reason for this is that non-native species tend to select different types of vegetation compared to native herbivores, leading to changes in plant community composition and microbial community composition. This change in vegetation reduces plant production and thus reduces carbon inputs to the soil. The lower relative abundance of *Rhizophagus* in the 2024 Bison and Grazer Areas may also indicate changes in plant– mycorrhizal interactions, as mycorrhizal fungi can contribute to plant nutrient acquisition and influence soil carbon dynamics. A study by Bagchi & Ritchie (2010) found that watersheds converted to livestock ranching had 49% less soil carbon compared to watersheds that retained native herbivores, despite similar grazing intensities and comparable animal densities. European bison are not known to have occurred naturally in Britain and should therefore be considered non-native in a strict biogeographical sense. Nevertheless, fossil and archaeogenomic evidence demonstrates that two other large wild bovids—the extinct steppe bison (*Bison priscus*) and aurochs (*Bos primigenius*)—formerly occurred in Britain (Gee, 1993; Park et al., 2015), and European bison are their closest living relative left. European bison may therefore represent a functional analogue of extinct British bovids rather than a direct replacement for a historically native species. Large herbivores were not present in West Blean and Thornden Woods prior to the introduction of bison. A common concern between many rewilding studies, is that there is a lack of understanding of the potential impacts of alternative species on the ecosystem, and thus unforeseen ecological risks (Fernández et al., 2017).

However, current evidence is not sufficient to demonstrate that soil carbon sequestration capacity is declining. The effects of stochastic variables such as climate change on the biomes in the study area are not negligible over the short time scale of two years. In other words, such short-term changes may not be a true reflection of long-term trends. Therefore, ongoing analyses should be conducted on longer time scales in the future. Furthermore, when analysing the quality of the extracted soil DNA, it was found that the DNA concentration in the 2024 Bison Area was much lower than other areas. Although normalization was done for the sequencing data before both relative abundance calculations and KEGG annotations, it is still possible that the data coverage of the 2024 Bison Area was incomplete. Finally, in addition to metagenomic analyses for biomes, more reliable conclusions need to be drawn in combination with specific soil physicochemical conditions and chemical data that more directly respond to soil carbon content such as total soil carbon, total nitrogen, and carbon to nitrogen ratios. In the future, analyses combining soil physicochemical data measured chemically or by quantitative PCR (qPCR) could better reveal the effects of European bison on soil biocomposition and function.

### 4.3 Functional shifts as early indicators of ecological change

Although no significant changes were detected in taxonomic alpha diversity, functional analyses revealed substantial differences in soil biotic functional composition. Functional beta diversity showed significant differences between the 2024 Bison Area and all three comparison groups. KEGG enrichment further identified consistent changes in carbon metabolism and amino acid biosynthesis. Therefore, in this study, functional shifts were detected, despite the absence of detectable changes in alpha diversity.

This suggests that the functional diversity and taxonomic diversity of soil biotic communities do not necessarily change in synchrony. Changes in metabolic activity of existing microbial taxa, or changes in the genetic potential for microbial metabolism may alter ecosystem functioning before pronounced changes in taxonomic diversity become evident. For example, Sveen et al. (2025) demonstrated that, during ecosystem development, the functional diversity of soil microbial communities can increase while taxonomic diversity decreases. Similarly, Louisson et al. (2023) showed that changes in the taxonomic composition and functional potential of soil microbial communities can become decoupled in response to land-use change. If functional changes can be detected despite limited changes in taxonomic alpha diversity, metagenomic approaches capable of recovering extensive functional information, including long-read approaches such as Oxford Nanopore sequencing, may therefore be useful for detecting subtle ecological change during early-stage of rewilding. This could have important implications for early-stage ecological monitoring of rewilding projects.

However, shotgun metagenomic data primarily reflect the presence and relative abundance of functional genes rather than their actual expression or metabolic activity. Therefore, the existing KEGG pathway abundance alone cannot demonstrate that a particular pathway is actively functioning or being expressed. Future transcriptomic analyses could provide a more direct assessment of the actual expression levels of different pathways, thereby offering a more accurate understanding of functional changes in soil microbial communities.

## Conclusion

Two years after the reintroduction of European bison, soil community alpha diversity remained unchanged, whereas significant shifts in community composition and functional profiles had begun to emerge. Consistent with this overall pattern, neither the alpha diversity of all detected soil taxa nor that of plant taxa changed significantly between 2022 and 2024. Nevertheless, relative-abundance analyses revealed emerging compositional changes, including decreases in several nitrogen-fixing bacterial genera and some mycorrhizal fungi. The difference in Beta diversity also demonstrates a change in the species structure of the 2024 Bison Area, and increased compositional heterogeneity. In terms of soil function, soil of 2024 Bison Area differs significantly from all the other comparing groups. The enrichment of amino acid biosynthesis and carbon metabolism pathways may be consistent with increased inputs of organic nitrogen and altered carbon processing. This may be due to the introduction of bison which has changed the local plant composition. Nevertheless, the available evidence is insufficient to determine whether soil carbon sequestration capacity has increased or decreased.

This study demonstrates the utility of Oxford Nanopore metagenomic sequencing for detecting taxonomic and functional variation in heterogeneous soil communities. Longer-term monitoring that integrates soil physicochemical properties, vegetation data, and other environmental variables will be required to distinguish bison-associated effects from temporal and spatial variation and to assess their consequences for soil functioning. Overall, these findings suggest that, during the early stages of temperate-forest rewilding, functional composition may provide a more sensitive indicator of ecological change than alpha-diversity metrics.

## Acknowledgments

We thank the Kent Wildlife Trust team for their support throughout this study, including facilitating site access and assisting with sample collection. We are especially grateful to Sally Smith for her outstanding contribution to project promotion.

We also thank the Genomics Core Laboratory at the MRC Laboratory of Medical Sciences, particularly Laurence Game and Ivan Andrew, for providing equipment support.

Special thanks go to the teams at the Natural History Museum: Fareeda Atwan and the Learning and National Programmes team, Evie Smith and the Special Events Development team, and Lottie Dodwell-Williams and the Public Space Programmes team, for their continued support, enthusiasm, promotion of the project, and for facilitating its inclusion in the *Fixing Our Broken Planet* Gallery.

